# Stromal LOX–FAK–β-catenin pathway locks mammary fibroblasts into a tumor-promoting myCAF state

**DOI:** 10.1101/2025.07.11.664481

**Authors:** Armin Gandhi, Divya Beri, Dharma Pally, Sathish Manjunath, Sharadhi Humcha, Annapoorni Rangarajan, Rekha V Kumar, Ramray Bhat, Utpal Tatu, Paturu Kondaiah

## Abstract

**Background:** Cancer-associated fibroblasts (CAFs) sustain tumor progression, yet the soluble cues that maintain their myofibroblast (myCAF) state are poorly defined. Transforming growth factor beta (TGF-β) is a canonical CAF activator. This study aims to identify TGF-β-induced secreted mediators that reinforce the myCAF phenotype in breast cancer and map the downstream signaling cascade.

**Methods and Results:** Secretome profiling of primary patient-derived myCAFs and human mammary fibroblasts (HMF3s) engineered to over-express TGF-β1 revealed 20 extracellular-matrix remodelers shared exclusively by both activated states; lysyl oxidase (LOX) was the top-ranked hit. LOX knockdown abrogated TGF-β–driven α-smooth-muscle actin (α-SMA) induction, collagen-gel contraction and migration in HMF3s, and reduced constitutive α-SMA and β-catenin in myCAFs. Mechanistically, TGF-β upregulated LOX, which activated focal-adhesion kinase (FAK), leading to p38 MAPK- and Akt-mediated Ser9 phosphorylation (inactivation) of GSK3β and consequent β-catenin stabilization. In HCC1806-luciferase orthotopic xenografts, CAFs accelerated tumor growth, whereas LOX-deficient CAFs lost this pro-tumoral effect.

**Conclusion:** LOX is a pivotal autocrine effector of TGF-β that locks breast CAFs into a pro-tumoral myCAF state through a LOX/FAK/GSK3β/β-catenin axis. Targeting stromal LOX may disrupt CAF activation and curb breast cancer progression.

## 1. Introduction

The tumor microenvironment (TME) is now recognized as a dynamic regulator of cancer initiation, progression and therapy resistance. Cancer-associated fibroblasts (CAFs) constitute a major stromal cell population that remodel the extracellular matrix (ECM), secrete growth factors and cytokines, and modulate immune cell infiltration, thereby facilitating invasion, metastasis and angiogenesis (1, 2). Recent single cell and spatial transcriptomic studies have underscored the heterogeneity of breast CAFs, revealing discrete myo-fibroblastic, inflammatory and antigen-presenting subsets with distinct functional programmes and clinical correlations (3, 4). Understanding how these phenotypes arise is therefore critical for therapeutic targeting.

Transforming growth factor-β (TGF-β) is a canonical inducer of myofibroblast differentiation marked by α-smooth muscle actin (α-SMA) expression and contractile cytoskeletal re-organization (5). Although α-SMA-positive “myCAFs” represent only one CAF subset, TGF-β signaling has emerged as a pivotal axis sustaining CAF activation across cancers (6). Beyond canonical SMAD-mediated transcription, TGF-β integrates with epigenetic modifiers such as GATA6 and TET1 to establish feed-forward loops that lock fibroblasts into a pro-tumoral state (7). In parallel, lysyl oxidase (LOX) family enzymes—upregulated in CAFs and secreted via small extracellular vesicles—catalyze ECM cross-linking, increase matrix stiffness and activate focal adhesion kinase (FAK)–YAP signaling to drive epithelial-mesenchymal transition (EMT) (8). LOX isoforms, including LOXL4, were recently shown to orchestrate annexin-mediated invasive machinery in triple-negative breast cancer, underscoring their multifaceted role in tumorigenesis (9). Despite these advances, the precise mechanisms by which TGF-β-induced LOX coordinates downstream signaling to maintain myCAF phenotype remain incompletely defined.

Here, we employed secretome profiling of α-SMA^+^ patient-derived breast CAFs and TGF-β1-overexpressing normal mammary fibroblasts to identify stromal factors that reinforce fibroblast activation. Comparative LC-ESI-Q-TOF-MS/MS revealed a set of 20 proteins shared exclusively between CAFs and TGF-β-stimulated fibroblasts, with LOX emerging as a leading candidate. Functional interrogation demonstrated that TGF-β-induced LOX activates a LOX/FAK/GSK3β/β-catenin cascade that is necessary for fibroblast contractility, migration and α-SMA induction. Finally, orthotopic xenograft experiments showed that LOX-deficient CAFs lose their tumor-promoting capacity, establishing LOX as a critical autocrine effector of TGF-β in the breast cancer stroma. Collectively, our study defines a previously unappreciated LOX-centered signaling hub that integrates mechanical and transcriptional cues to stabilize the myCAF state and highlights stromal LOX as an actionable target to disrupt CAF-mediated breast cancer progression.

## 2. Materials and Methods

### 2.1 Cell lines and treatments

HMF3s, human mammary fibroblasts immortalized with human TERT and SV40 large T antigen (10) were a generous gift from Dr. P. Jat. Cancer associated fibroblasts (CAFs) were isolated from human breast cancer biopsy tissues obtained from patients treated at Kidwai Memorial Institute of Oncology as previously described (11). Experiments with patient samples were carried out after approval from the Institutional Ethical Committee. All experiments with patient derived CAFs were completed within the 12^th^ passage of the cells (12). Fibroblasts were cultured at 37 °C, in a humidified chamber with 5 % CO_2_ in growth media consisting of DMEM (Sigma Aldrich, USA) supplemented with 10 % Newborn Calf Serum (NBCS). HCC1806 cells were cultured in DMEM (Sigma-Aldrich, USA) medium supplemented with 10 % foetal bovine serum (FBS). All cell cultures were supplemented with an antibiotic cocktail of penicillin (100 units/ml) and streptomycin (100 ug/ml) (Invitrogen Life Sciences, USA). FBS was heat inactivated at 55 °C for 30 minutes prior to use. All the treatments with rhTGF-β1 and inhibitors was in serum free conditions. Cells were allowed to grow till 80 % confluence and cultured in serum free media for 24 hours prior to treatments. The following inhibitors were utilized: β-catenin/TCF inhibitor (FH535, 10 µM; Sigma Aldrich, USA); FAK inhibitor (FAK inhibitor I, 10 µM; MERCK, USA); PI3K inhibitor (LY249002, 10 μM; CST, USA); p38 inhibitor (SB203580, 10 μM; CST, USA); Erk inhibitor (PD98059, 10 μM; Sigma Aldrich, USA); JNK inhibitor (SP600129, 10 μM; Calbiochem, USA); LiCl (20 mM; Sigma Aldrich, USA). The inhibitor treatments were done 2 hours prior to rhTGF-β1 (5 ng/ml; R&D systems, USA) stimulation.

### 2.2 Reagents and antibodies

Anti phospho Smad3 (Ser423/425, ab52903) and anti α-SMA (ab5694) antibodies were purchased from Abcam, USA. Anti phospho GSK3β (Ser9, #9323), GSK3β (#9315), phospho FAK (Tyr397, #3283S), FAK (#3285S), Smad3 (#9523), phospho Akt (Ser473, #9271), Akt (#9272), p38 (#9212S), β-actin HRP (#5125) and α-tubulin HRP (#9099S) were purchased from Cell Signaling Technologies, USA. Anti-LOX antibody (NBP2-24877) was purchased from Novus Biologicals, USA. Anti phospho p38 (Thr180/182, IMG-679) was purchased from Imgenex, India. Anti β-catenin antibody (C2206), and Vimentin (V2258) were purchased from Sigma Aldrich, USA. HRP-conjugated goat anti rabbit IgG (HPO3) was purchased from Geneilabs. Anti-rabbit immunoglobulin conjugated with Alexa Fluor 488 was purchased from Molecular Probes, ThermoFischer Scientific, USA.

### 2.3 TGF-β1 over-expression in HMF3s cells

Mutant porcine TGF-β1 cDNA (Ser 223, 225) isolated from pPK9a construct (13) was subcloned into pBABEPuro vector (pBPTGF-β1). This cDNA construct expresses activated form of TGF-β1. Note, mature porcine TGF-β1 is identical to the human TGF-β1 (14). The pBPTGF-β1 was retrovirally transduced into HMF3s cells for stable expression of TGF-β1 in fibroblast cells. For virus generation, 70% confluent HEK293t cells in 60 mm dishes were transfected using Lipofectamine 3000 reagent with 1 μg pBABEPuro or pBABEPuroTGF-β1 along with retroviral helper plasmids, pUMVC3 (packaging plasmid, 800 ng) and pCMV-VSV-G (envelope plasmid, 200 ng). After 24 hours of recovery in DMEM with 10 % FBS, conditioned media containing viral particles were collected over a period of 48 hours and stored at ™80°C. HMF3s cells were cultured to 80 % confluence in 60 mm dishes. The virus containing conditioned medium was thawed on ice. Protamine sulphate at a final concentration of 10 μg/ml was added to 3 ml of conditioned medium. The pH of the medium was maintained at 7.2 by addition of HEPES buffer solution. The media was then passed through a 0.45 μm filter and added onto the HMF3s cells. The cells were incubated with the virus for 12 hours at 37°C in CO_2_ incubator. After 12 hours, the media was replaced with fresh growth media. After 24-48 hours the cells were transferred to 90 mm dishes and allowed to attain confluence. The over-expressing cells were selected with 1 μg/ml puromycin for 7-10 days. Cells were maintained in culture with 0.2 μg/ml puromycin. TGF-β1 over-expression was confirmed by transcript analysis by semi-quantitative PCR and PAI-luciferase reporter assay.

### 2.4 AI-luciferase reporter assay

To detect active TGF-β in the conditioned media of HMF3s cells, CCL64 cells expressing luciferase fused to PAI promoter (CCL64PAI cells) (15) were used as described previously (16). Briefly, CCL64PAI cells were plated in 96 well plates and treated with 24-hour conditioned media from HMF3s cells. Cells were lysed and luciferase activity was measured using luciferase assay kit (Promega, USA) as per manufacturer’s protocol in a luminometer (TD-20/20 Turner Designs, USA). The reading obtained as Relative Luciferase Units (RLU) was plotted.

### 2.5 Transient transfection and knockdown of genes

Transient transfection of plasmids was performed using Lipofectamine 2000 reagent (Thermo Scientific, USA) as per manufacturer’s instructions. LOX and β-catenin shRNA constructs were procured from MISSION pKLO.1 shRNA library, Sigma Aldrich (A kind gift from Prof. G. Subbarao).

### 2.6 TOP-FLASH assay

TOP-FLASH assay was performed to study the transcriptional activity of β-catenin. Briefly, 2 × 10^5^ were plated in 24-well dishes and co-transfected with pSuperTOP-FLASH-lux (Upstate Cell Signaling Solutions, USA) and pRL-TK (renilla luciferase, for normalization) constructs at 100:1 ratio respectively using lipofectamine 3000 (Invitrogen, USA) reagent as per the manufacturer’s protocol. pFOPFLASH-lux (Upstate Cell Signaling Solutions, USA) with mutated β-catenin binding sites was used as a negative control. Following transfection, cells were serum starved and treated with TGF-β at 48 hours. Cells were lysed in passive lysis buffer and the lysates were processed for luciferase activities (Dual-Luciferase Reporter Assay System, Promega, USA). The ratio of firefly to renilla readings were obtained in a luminometer (TD-20/20, Turner Designs, USA) and the fold change with respect to the untreated control was plotted.

### 2.7 RNA isolation and quantitative PCR analyses

Total RNA was isolated from cells using TRI reagent (Sigma-Aldrich, USA) as per manufacturer’s protocol. The RNA quantity and quality were assessed by OD_260_ and OD_280_ measurements on a spectrophotometer (Nanodrop ND1000, Thermo-scientific, USA) and the integrity was determined on a MOPS-formaldehyde denaturing agarose gel. For cDNA synthesis, two micrograms of total RNA was reverse transcribed using the ABI cDNA High-Capacity cDNA synthesis kit (Applied Biosystems, USA). PCR reactions were done using DynamoSYBR green mix (Finnzymes, Finland) in ViiA 7 Real-Time PCR system (Applied Biosystems, USA). The analysis was done using QuantStudio software (Applied Biosystems, USA). RPL-35A expression was used for normalization. Gene specific primers used for amplification and detection of gene products have been listed in table S1.

### 2.8 Western blot analysis

Cells were washed with chilled phosphate buffered saline, pH 7.4 (PBS) and lysed with RIPA buffer (50 mM Tris–Cl pH-8.0; 150 mM NaCl; 0.1 % SDS; 1.0 % NP40; 0.5 % sodium deoxycholate; protease inhibitor cocktail, PIC; 1 mM PMSF; 5 mM EDTA; 10 mM sodium fluoride and 1 mM sodium orthovanadate) with constant agitation at 4^°^C for 30 minutes. Cell debris were removed by centrifugation at 12,000 rpm for 20 minutes. Total protein content in the cell lysates was estimated by Bradford assay (Biorad, USA). Equal amount of protein was loaded onto 10 or 12.5% SDS-PAGE gels and transferred onto PVDF membranes (ImmobilonP, Millipore Corporation, Germany) by using wet electrophoresis transfer apparatus (Biorad, USA). Blots were blocked for an hour at room temperature with 5% non-fat dry milk (Sigma-Aldrich, USA) made in TBST buffer (20 mM Tris-HCl pH 7.4, 137 mM NaCl, 0.1% Tween-20) and probed overnight with primary antibodies (1:1000) at 4^°^C. After washing off unbound antibody, blots were incubated with HRP conjugated anti Rabbit IgG for 45-60 minutes (1:5000) at room temperature. Membranes were probed with femtoLUCENT PLUS-HRP chemiluminescent reagent (G-Biosciences, USA). Signal was captured using gel documentation system (Biorad, USA).

### 2.9 Immunocytochemistry

Approximately 50,000 cells were plated on sterile cover slips. After treatments, cells were fixed in 3.7 % paraformaldehyde for 15 minutes at room temperature, washed with phosphate buffered saline (PBS, pH 7.4) and permeabilized with chilled methanol for 10 minutes at – 20 ^°^C. Non-specific binding of antibody was blocked by 5 % BSA and 0.03 % Triton X 100 in PBS for 1 hour at room temperature. Cover slips were incubated with primary antibody (1:100) overnight at 4^°^C, rinsed in PBS and probed with Alexa fluor tagged antiRabbit IgG secondary antibody (1:250) for 1 hour at room temperature. Following PBS washes, nuclei were stained with DAPI and mounted with ProLong Gold antifade reagent (Life technologies, USA) on clean glass slides. The fluorescence was visualized and documented using Leica TCS SP5 confocal microscope.

### 2.10 Collagen contraction assay

Collagen contraction assay was performed to assess the contractile phenotype of fibroblasts, as previously described (17). Briefly, 4 × 10^5^ cells were suspended in 660 µl serum free DMEM. 330 µl of chilled rat tail collagen (3 mg/ml in 20 mM acetic acid solution, Thermo Fisher Scientific, USA) was added to the cell suspension and the pH was immediately adjusted to 7.2 with 1N NaOH. The mixture was then gently poured in 12 well plates and allowed to set at room temperature for 20 minutes followed by incubation at 37 ^°^C for 40 minutes. After the gel is firm, serum free DMEM was gently poured into the wells and the gels were detached by scooping with P200 tip and gentle tapping of the dish. The gels were imaged after treatments at 24-hour, 48-hour and 72-hour using gel documentation system (Biorad, USA). The images were analysed in ImageJ and the percent contraction was calculated.

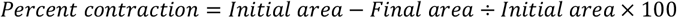

### 2.11 Wound healing assay

HMF3s cells were plated in 6 well plates and serum starved for 24 hours. A scratch was created by making two scratches perpendicular to each other using a P200 tip. Cells were washed and TGF-β treatment was initiated. Images were captured at 0-hour and 12-hour time points in Carl Zeiss microscope using Zeiss Zen software. Quantification of wound closure was done with wavelet transformation analysis in MATLAB.

### 2.12 Mass spectrometry

Conditioned medium (40 ml) was concentrated by ultrafiltration to less than 200 µl and analysed using Bradford reagent. 100 µg of total protein was subjected to in-solution trypsin digestion. The LC MS/MS and data analysis was performed as described previously (18). Briefly, 100 ug of proteins were reduced using DTT (10 mM), alkylated with iodoacetamide (20 mM) and subjected to in-solution trypsin (Proteomics grade, Promega Gold, 1 µg/µl) digestion at 37^°^C for 16 hours at pH 8.0. Trypsin was inactivated with formic acid, samples were dried under vacuum and resuspended in MS grade water prior to liquid chromatography/tandem mass spectrometry (LC-MS/MS) analysis. A blank control with plain media was run similarly. The raw data were analysed using Protein Pilot. Peptide scores above 50 and a protein score of minimum 1.3 corresponding to a confidence level greater than 95 % were used. The mass spectrometry proteomics data have been deposited to the ProteomeXchange Consortium (19) through the PRIDE (20) partner repository with the dataset identifier PXD022374 and 10.6019/PXD022374. (Reviewer Account details: Username; password: Wzzqlet). The data will be made public upon publication.

### 2.13 In vivo experiments

All experimental protocols using animals were approved by the Institutional Animal Ethics Committee (IAEC) of Indian Institute of Science, Bangalore. Female nude mice: 4-6 weeks of age were housed in clean air facility. Autoclaved water and mouse chow were provided *ad libitum*. The bedding and water were changed every 4 days. HCC1806 cells were transfected with pLentiluc construct (Lipofectamine 3000) and selected with 1 ug/ml puromycin to obtain stable expression. The luminescence was confirmed in vitro using Luciferase assay kit (Promega) following manufacturer’s protocol. For assessing tumorigenesis, luciferase expressing HCC1806 cells were mixed with fibroblasts and injected into the mammary fat pad of 4-6 weeks old female Balb/c nude mice. Cancer cells were trypsinized and counted to a suspension of 10^7^ cells/ml. HMF3s or CAFs were trypsinized and counted to make a suspension of 2×10^6^ cells/ml (20 % of cancer cells). Suspension of cancer cells was thoroughly mixed with fibroblast cells. The mice were anaesthetized with a solution of ketamine and xylazine (50 mg/kg ketamine and 10 mg/kg xylazine). The mammary fat pad was located surgically, 100 μL cell suspension was injected and the incision was sutured. The tumour growth was monitored with *in vivo* bioluminescence imaging and tumour diameter measurements. Tumour diameter was measured with vernier callipers and tumour volume was calculated using the formula,

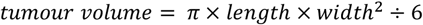

Where, length represents the largest tumour diameter and width represents the perpendicular tumour diameter. For bioluminescence imaging, animals were injected with 100 μl of D-luciferin intraperitoneally (15 mg/ml, Promega) and anaesthetized using isoflurane saturated with oxygen. The cells were imaged using IVIS (*In Vivo* Imaging System) Platform (PerkinElmer, U. K.) within 5 minutes of injection. An overlap of the cells on the host was generated by the imaging software. The images were analysed using LivingImage software.

### 2.14 Statistical analysis

To compare two groups, Student’s *t* test was performed with 95% confidence interval. When comparing more than one group with or without inhibitor or gene knock down, two-way analysis of variance (ANOVA) followed by Bonferroni post-tests with confidence interval of 95% was performed for obtaining p values. P value ≤ 0.05 was considered significant. Numerical data are expressed as mean ± standard deviation. Graph Pad Prism 5 statistical package was used for statistical analysis and representation of numerical data. *P <0.05; **P<0.01; ***P<0.001.

## 3. Results

### 3.1 Secretome profiling reveals LOX as a critical mediator of TGF-β–driven fibroblast activation

CAFs are thought to arise predominantly from activation of resident fibroblasts within the tumour microenvironment (12, 21, 22). TGF-β–SMAD signalling drives fibroblast activation in fibrosis (23, 24) and in cancer (25-27). We therefore interrogated the TGF-β-responsive secretome for factors that might stabilise the CAF phenotype in breast cancer. Patient-derived CAFs (CAF-1 to CAF-3) were expanded in vitro; all cultures stained positive for α-SMA and vimentin, confirming a myCAF phenotype (Fig. S1A). Conditioned media (48 h) from these CAFs and from HMF3s fibroblasts ± TGF-β1 over-expression (Fig. S1B, S1C confirming over-expression by RNA and luciferase assay respectively) were analysed by LC-MS/MS. Unique-peptide filtering (≥2 peptides) identified 441, 470 and 1 055 proteins in CAF-1, CAF-2 and CAF-3 secretomes respectively. A Venn comparison revealed ∼185 proteins common to all fibroblast states, 12 shared only between HMF3s and CAF-1, and 20 shared only between CAF-1 and TGF-β-expressing HMF3s (Fig. 1A). Similar overlap patterns were obtained for CAF-2 and CAF-3 (Fig. S1D, S1E), attesting to dataset robustness. Reactome analyses (STRING database) highlighted enrichment of ECM structural/remodelling proteins (Fig. 1B, S1F and S1G). LOX emerged as a common hit among all the CAF- and TGF-β–shared proteins (Fig. 1A, S1B, S1C). LC-MS/MS hits including COL5A1, IGFBP3, LTBP1, LTBP2, LOX, SERPIN1, TSP1, VCAN were validated in HMF3s overexpressing TGF beta, in HMF3s treated with ectopic human recombinant TGF-β and additional breast CAFs from human patients by RNA expression analysis (Fig S1H, S1I, S1J). Because LOX family enzymes have been implicated in desmoplasia (28, 29) and tumour progression (30, 31) we hypothesised that LOX might be required for TGF-β– driven fibroblast activation. Transient LOX knock-down (shRNAs G3 and G4) in HMF3s fibroblasts reduced TGF-β–induced α-SMA protein to baseline (Fig. 1C), indicating that LOX is necessary for myofibroblastic differentiation. To test functional consequences, LOX-deficient cells were subjected to collagen-gel contraction and wound-healing assays. LOX knock-down almost completely abrogated TGF-β–induced gel contraction (Fig. 1D) and reduced wound closure from ∼45 % to <15 % after 12 h (Fig. 1E). These findings collectively establish LOX as a key autocrine effector of TGF-β in mammary fibroblasts.

**Figure 1.**
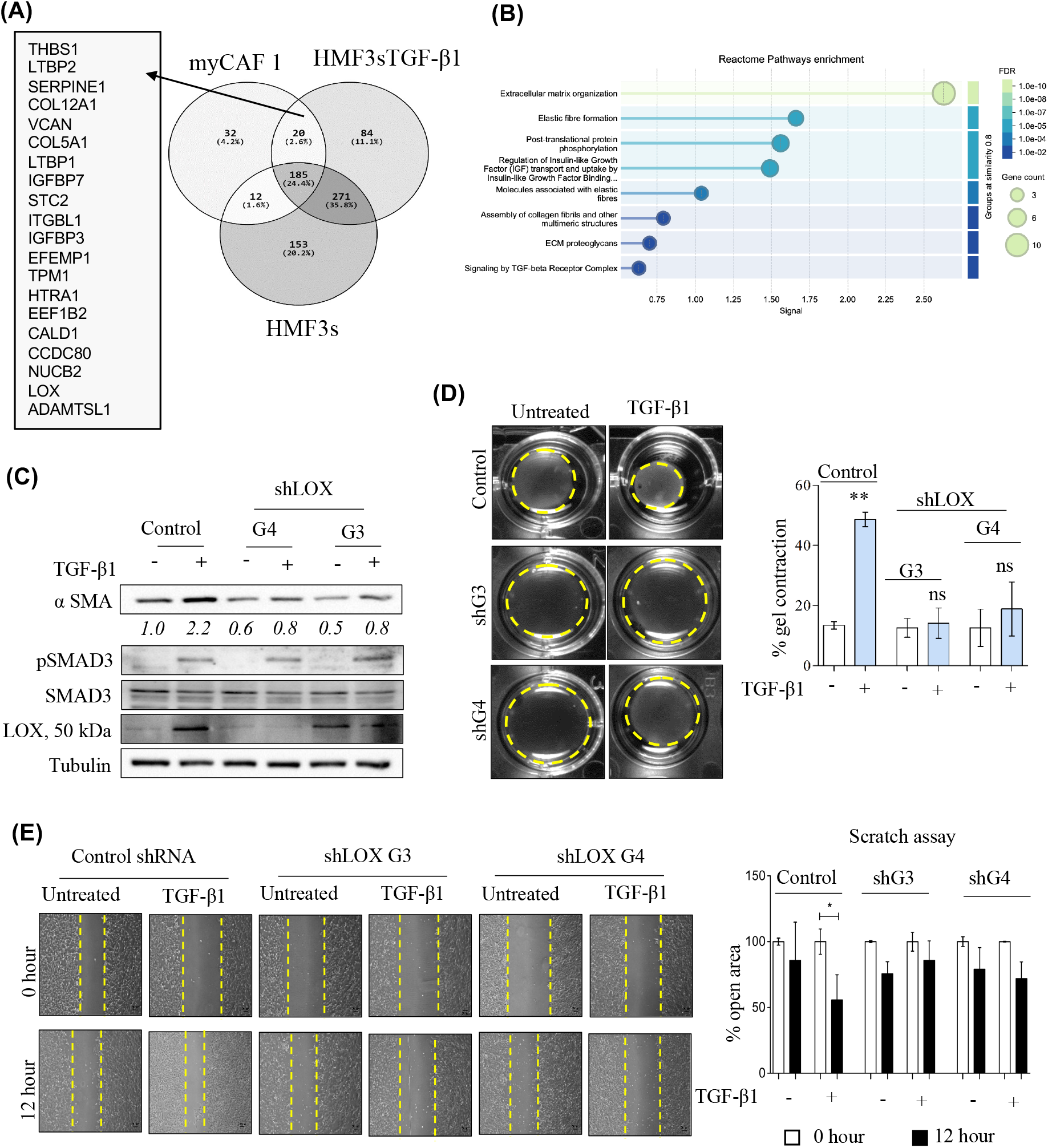
Secretome profiling reveals LOX as a critical mediator of TGF-β–driven fibroblast activation. **(A)** Representative Venn diagram of proteins secreted by myCAF-1 and HMF3sTGF-β cells compared to normal fibroblasts (HMF3s). 20 proteins are exclusively common to HMF3sTGF-β and myCAF-1. **(B)** Plot showing Reactome pathway analysis of 20 shared proteins common to HMF3sTGF-β and myCAF-1. **(C)** Western blot showing expression of α-SMA, pSmad3 and LOX upon TGF-β treatment in HMF3s cells transiently transfected with shRNA (G3 and G4) against LOX. Total Smad3 and Tubulin are used as internal controls. The band for LOX is at the apparent molecular weight of 50 kDa, the expected molecular weight of the LOX proenzyme. Numbers below the blot represent fold change in density values from multiple experiments normalized to α-tubulin with respect to untreated controls. **(D)** Collagen gel contraction assay of HMF3s cells in LOX knock down cells upon TGF-β treatment. Collagen matrix and contraction of the gel was monitored for a period of 48-hours and percent gel contraction quantitated and plotted. **(E)** Wound healing assay (Scratch Assay) using HMF3s cells treated with TGF-β and LOX knockdown. Photographs show visual representation of the wound closure and the bar diagram depicts quantitation of the wound closure. N = 3 for all the experiments; *P value < 0.05; ** P value < 0.01, Scale bar 20 µm, magnification 10X.

### 3.2 TGF-β–induced α-SMA depends on LOX-mediated FAK activation and β-catenin stabilisation

Previous studies have shown that LOX activates FAK through phosphorylation. FAK activation has been shown to cause fibroblast transition to myofibroblasts. We next hypothesized that LOX mediates alpha SMA induction and myofibroblast activation through FAK activation. Pharmacological FAK blockade (FAK inhibitor-1) abrogated TGF-β–driven α-SMA induction (Fig. 2A), while LOX knock-down suppressed both basal and TGF-β–stimulated FAK phosphorylation (Fig. 2B). These data position LOX upstream of FAK in the fibroblast-activation cascade. Previous studies implicated β-catenin signalling in TGF-β–mediated fibroblast activation (32, 33). Consistently, we observe translocation of beta catenin to the nucleus and upregulation of beta catenin activity upon TGF-β1 treatment onHMF3s cells (Fig. 2C, 2D). In addition, β-catenin/TCF blockade or shRNA-mediated knockdown reduced TGF-β–induced ACTA2/α-SMA mRNA from a 1.98-to a 1.13-fold change (Fig. 2E, 2G), diminished α-SMA protein in both immunoblot and immunocytochemistry assays (Fig. 2F, 2H, 2I, S2A) and compromised contractile ability of fibroblasts collagen gels (Fig. 2J). In addition, LOX silencing and FAK inhibition curtailed GSK3β-Ser9 phosphorylation (Fig. 3K, 3L) linking the LOX-FAK axis directly to canonical β-catenin activation. Notably, Wnt2 and Wnt4 expression remained unchanged (Fig. S2B), excluding autocrine Wnt ligands as alternative β-catenin activators. Together with our data, these findings consolidate LOX-mediated FAK activation as a pivotal driver of β-catenin–dependent fibroblast activation. In summary, TGF-β-induced LOX activates FAK, leading to GSK3β inhibition, β-catenin stabilisation and α-SMA up-regulation—thereby locking mammary fibroblasts into a myofibroblast state.

**Figure 2.**
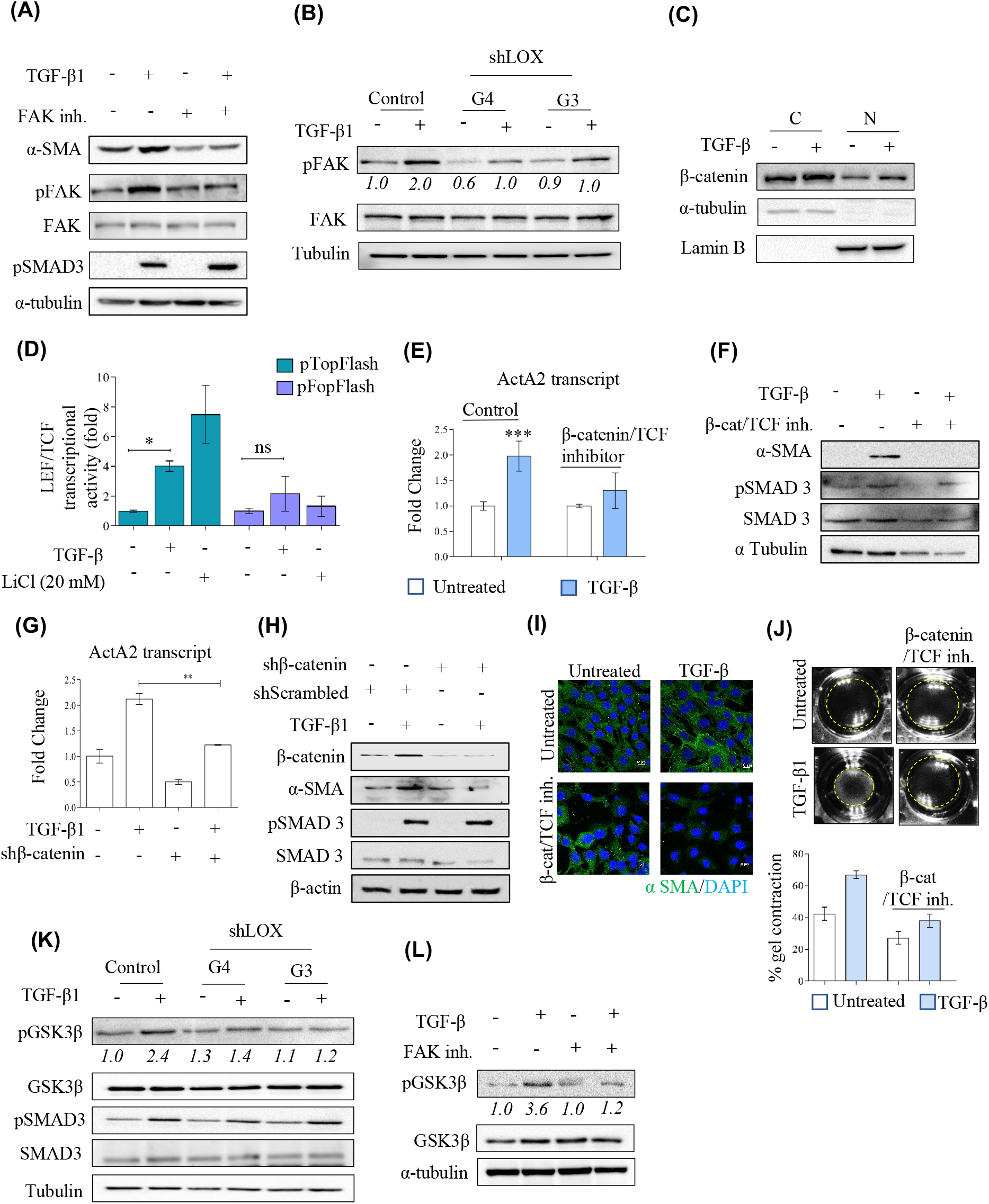
TGF-β–induced α-SMA depends on LOX-mediated FAK activation and β-catenin stabilisation. **(A)** Western blot analysis of α-SMA and pSmad3 upon TGF-β treatment and/or FAK inhibition. Expression of TGF-β induced expression of α-SMA is compromised upon FAK inhibitor treatment. FAK and α-tubulin are internal controls. **(B)** Western blot analysis showing pFAK and pSmad3 upon TGF-β treatment and/or LOX knockdown. Note the compromised TGF-β–induced phosphorylation of FAK upon LOX knockdown using LOX shG3 and shG4. Numbers below the blot indicate fold change in densitometry values normalized to α-tubulin and with respect to untreated controls from three different experiments. **(C)** Western blot analysis of β-catenin in the nuclear (N) and cytoplasmic (C) fraction of HMF3s cells with and without TGF-β treatment. α-tubulin is the internal control for cytoplasmic fraction and Lamin B1 is the internal control for nuclear fraction. **(D)** Graph depicts fold change of luciferase activity of pTOPFLASH indicative of β-catenin activity. TGF-β increased β-catenin activity by approximately 4.5 folds. LiCl (20 µM) was used as positive control respectively. pFOPFLASH plasmid consisting of a mutated Tcf/Lef binding promoter was used as the negative control for pTOPFLASH. **(E)** Graph depicting fold change of ActA2 transcript in HMF3s cells in the presence of TGF-β and/or small molecule inhibitor for β-catenin. HMF3s cells were treated with β-catenin/TCF inhibitor and analysed for ActA2 transcript and protein expression. β-catenin/TCF inhibitor compromised TGF-β–induced ActA2 transcript in HMF3s cells. **(F)** Western blot analysis of α-SMA and pSmad3 in HMF3s cells in the presence of TGF-β and/or β-catenin/TCF inhibitor. Presence of β-catenin/TCF inhibitor compromised TGF-β–induced α-SMA protein. **(G)** Graph depicts fold change of ACTA2 in HMF3s cells upon TGF-β treatment with respect to untreated controls. TGF-β–induced ActA2 transcript was compromised in the presence of β-catenin shRNA in HMF3s cells. **(H)** Western blot analysis of α-SMA, β-catenin and pSmad3 in HMF3s cells in the presence of TGF-β and/or a transient knockdown of β-catenin. Knockdown of β-catenin in HMF3s compromised TGF-β–induced α-SMA expression. **(I)** Representative images showing immunocytochemical analysis of TGF-β–induced α-SMA in HMF3s cells in the presence of β-catenin/TCF inhibitor. Presence of β-catenin/TCF inhibitor compromised TGF-β–induced α-SMA in HMF3s cells. Scale bar 10 µm, magnification 63×. **(J)** Representative images of collagen gels embedded with HMF3s cells. Graph depicts percent gel contraction calculated from three independent experiments. TGF-β induced collagen gel contraction was compromised in the presence of β-catenin/TCF inhibitors. **(K)** Western blot analysis of pGSK3β in HMF3s cells upon TGF-β treatment and with transient transfection of shRNA against LOX. TGF-β– induced pGSK3β were compromised upon LOX knockdown with shG3 and shG4 transfection. **(L)** Western blot depicting TGF-β–induced pGSK3β in HMF3s cells upon FAK inhibition. FAK inhibitor abrogated TGF-β–induced pGSK3β. GSK3β and α-tubulin are internal controls. N = 3; *P value < 0.05; ** P value < 0.01 ***P value < 0.001

**Figure 3.**
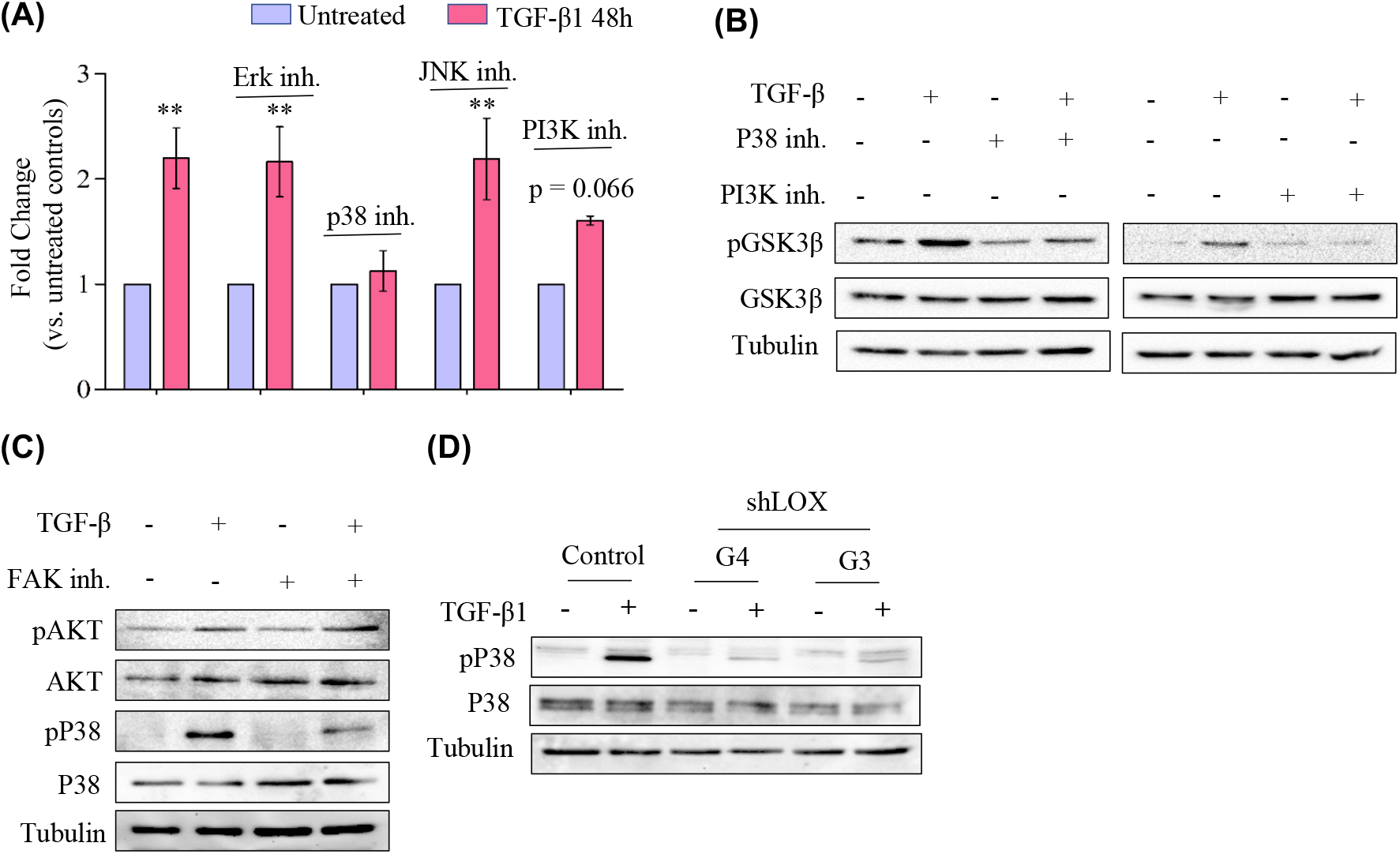
p38 MAPK and Akt cooperatively inhibit GSK3β to sustain TGF-β–driven fibroblast activation. **(A)** Graph depicts fold change for ActA2 in HMF3s cells upon TGF-β treatment with or without Erk, p38, JNK and PI3K inhibitors respectively. Presence of p38 MAPK and PI3K inhibitors compromised TGF-β induced ActA2 transcript in HMF3s cells. **(B)** Western blot analysis of phospho GSK3β upon TGF-β treatment in the presence or absence of p38 MAPK and PI3K inhibitors. Inhibition of both p38 MAPK and PI3K compromised TGF-β induced phosphorylation of GSK3β at serine 9. **(C)** Western blot analysis of phospho p38 and phospho Akt upon TGF-β treatment with or without FAK inhibition. FAK inhibition compromised TGF-β induced phospho p38 and had no effect on phospho Akt in HMF3s cells. **(D)** Western blot analysis of phospho p38 and p38 in the presence of TGF-β and with or without LOX knockdown with shRNAs G3 and G4. TGF-β induced phospho p38 was compromised upon LOX knockdown in HMF3s cells.

### 3.3. p38 MAPK and Akt cooperatively inhibit GSK3β to sustain TGF-β–driven fibroblast activation

Our previous results positioned LOX and FAK upstream of GSK3β inhibition; here we dissect the kinase hierarchy that inactivates GSK3β (Ser9) and thereby stabilises β-catenin during TGF-β signalling. Protein kinase B/Akt is a well-established Ser9 kinase (34) and MAPKs have also been implicated (34-36). To test their relative contributions, HMF3s fibroblasts were stimulated with TGF-β in the presence of selective PI3K (Akt pathway) or MAPK inhibitors (ERK, JNK, p38). TGF-β elevated ACTA2 mRNA 2.2-fold; ERK or JNK blockade left this induction unchanged (2.16- and 2.19-fold, respectively), whereas p38 and PI3K inhibition reduced it to 1.13- and 1.60-fold (Fig. 3A). Thus, p38 MAPK and Akt, but not ERK/JNK, contribute to ACTA2 transcription. Immunoblotting confirmed that p38 or PI3K inhibition attenuated TGF-β-induced GSK3β-Ser9 phosphorylation (Fig. 3B), aligning biochemical read-outs with the transcript data. Because LOX and FAK lie upstream, we next asked whether they modulate p38 or Akt. FAK inhibition suppressed p38 phosphorylation without affecting Akt (Fig. 3C), and LOX knock-down likewise diminished p38 activation (Fig. 3D). These findings place p38 downstream of the LOX/FAK axis, whereas Akt activity appears independent of LOX/FAK but still necessary for full GSK3β inhibition. Collectively, our data support a pathway in which TGF-β signals through LOX, FAK and p38 and in parallel Akt inputs to phosphorylate GSK3β (Ser9), stabilise β-catenin and drive α-SMA expression.

### 3.4 LOX sustains the myCAF phenotype and fuels CAF-driven mammary-tumour growth in vivo

Based on our previous observations, we hypothesized that LOX is constitutively expressed in patient-derived myCAFs and is required to maintain their α-SMA^+^/β-catenin^+^ state. In line with these observations, stable LOX knock-down in three independent patient-derived breast-CAFs (CAF-1a, CAF-4a, CAF-5a) reduced α-SMA protein by ∼50 % and β-catenin by 40–80 % (Fig. 4A and S4A). These data indicate that ongoing LOX expression is essential to lock CAFs into an activated, myofibroblastic phenotype, consistent with recent reports linking stromal LOX to CAF contractility and YAP/β-catenin co-signalling (37). Clinical datasets corroborate our in-vitro findings. In a public breast-cancer RNA-seq cohort (38), LOX mRNA was significantly higher in bulk tumours than in normal breast (P = 0.0049; Fig. 4B). Oncomine meta-analysis (https://www.oncomine.org/resource/login.html) further showed a stromal-specific LOX up-regulation in both invasive ductal carcinoma (IDC; median ×3.7, P = 6.1 × 10^−4^) and ductal carcinoma in situ (DCIS; median ×1.8, P = 0.001), whereas epithelial LOX levels remained unchanged (Fig. 4C-4F). These outcomes fit with recent spatial-omics work demonstrating that breast-tumour stiffness and LOX-mediated collagen cross-linking arise largely from CAFs rather than carcinoma cells (39). To text if LOX-deficient CAFs lose their tumour-promoting capacity, we performed orthotopic co-injection of HCC1806-luc breast-cancer cell line with CAF-5a or CAF-4a. Co-inoculation of HCC1806 with CAF-4a and CAF-5a accelerated tumour growth, increasing volume and bioluminescence relative to HCC1806 ± normal fibroblasts (HMF3s). LOX knock-down in either CAF line significantly blunted this effect: tumour volume dropped >40 % and photon flux fell by ∼45 % at 4 weeks (Fig. 4G, 4H, S3B, S3C). The modest residual growth advantage over HMF3s likely reflects incomplete LOX suppression. These results mirror recent studies in pancreatic and oral carcinomas where pharmacological LOX inhibition decreased stromal stiffness and curtailed tumour expansion (8, 40). Together, our data establish LOX as a dual-function effector: autocrine maintenance of the myCAF α-SMA/β-catenin programme and paracrine enhancement of breast-tumour growth. Given the emergence of clinically tractable pan-LOX inhibitors, our findings provide a strong rationale for testing stromal-targeted LOX blockade as an adjuvant therapy in aggressive, desmoplastic breast cancers.

**Figure 4.**
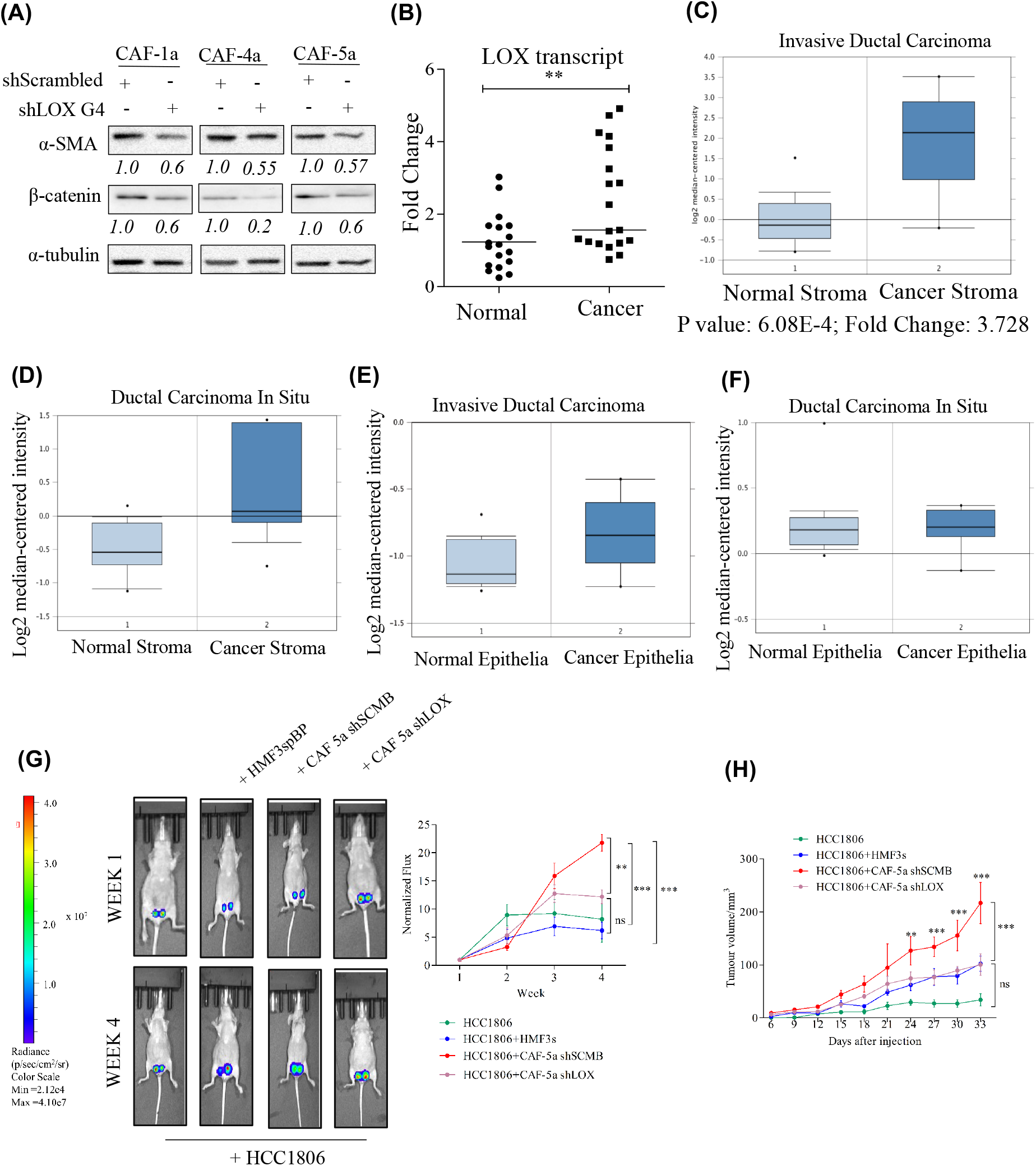
LOX sustains the myCAF phenotype and fuels CAF-driven mammary-tumour growth in vivo. (**A**) LOX regulates α-SMA expression and total β-catenin in myCAFs. Western blot analysis of α-SMA and β-catenin expression in three myCAFs isolated from three different breast cancer biopsy tissue samples. Stable knockdown of LOX in myCAFs downregulated α-SMA and β-catenin as compared to scrambled control. α-tubulin is the internal control. Numbers below the blots represent fold change with respect to untreated controls calculated from densitometry analyses normalized with α-tubulin from at least three independent experiments. LOX expression is upregulated in breast cancer tissues. A scatter plot of Real time PCR analysis of LOX in whole breast cancer patient tissues with respect to normal samples. Horizontal line in the scatter represents median, ** p value < 0.005. LOX transcript is upregulated in majority of breast cancer biopsy samples. **(C), (D)**. Oncomine database analysis of LOX in stroma of IDC and DCIS respectively [GEO: GSE14548]. Data shows upregulation of LOX in breast cancer stroma as compared to normal stroma. LOX is upregulated in IDC stroma with median fold change of 3.728 (p = 6.08E-4) and in the stroma of DCIS samples by median fold change of 1.848 (p = 0.001) with respect to normal stroma. **(E), (F)**. Oncomine database analysis of LOX in epithelia of IDC and DCIS respectively [GEO: GSE14548]. LOX is upregulated by median fold change of 1.153 (p = 0.027) in IDC samples and downregulated by median fold change of 1.019 (p value 0.629) in DCIS patient samples in comparison to normal breast tissue samples. **(G)** Representative images of bioluminescence imaging of HCC1806 tumours in female immunocompromised mice. The plot represents photon flux of HCC1806 tumours measured weekly normalized to the signal obtained at the first week of injection. HCC1806 showed significantly increased growth when injected with CAF-5a cells. LOX knockdown in CAF-5a compromised increased bioluminescence of HCC1806 cells. N = 6, *p value < 0.05, **p value < 0.01, ***p value < 0.005. **(H)** The plot represents HCC1806 tumor volume with respect to time. Tumor volumes are significantly higher for HCC1806 injected with CAF-5a scrambled control as compared to those implanted alone or with HMF3s (vector control) or with LOX deficient CAF-5a.

## 4. Discussion

CAFs originate from multiple sources—including resident fibroblasts, bone-marrow–derived mesenchymal stromal cells, pericytes and adipocyte progenitors—as shown by single cell studies in breast cancer (4, 39). Their phenotypic diversity is now commonly resolved into at least three major subsets: inflammatory CAFs (iCAFs), myofibroblastic CAFs (myCAFs) and antigen-presenting CAFs (apCAFs) (4, 6, 41). Emerging sub-clusters such as “metabolic” and “vascular-associated” CAFs have also been described in spatial-omics atlases (42, 43). Up-regulation of TGF-β and its transcriptional targets is a shared feature of desmoplastic niches (44) and TGF-β signaling is increasingly recognized as a master switch for myCAF differentiation (45, 46). We therefore set out to delineate how TGF-β reprograms mammary fibroblasts into a stable myCAF state and how this, in turn, supports tumor progression. Secretome profiling uncovered 20 TGF-β–shared proteins, most of which are ECM components or ECM-remodeling enzymes—a hallmark of the myCAF proteome (6) that correlates with poor outcome (47). LOX emerged as a one of the common targets shared between CAFs and TGF-β1 over-expressing human mammary fibroblast cell line. Whereas LOX has long been linked to cancer cell invasion (48-50), our data identify LOX as a stromal effector that is necessary for TGF-β–driven fibroblast activation, encompassing α-SMA expression, collagen gel contraction and migration. These findings complement studies showing that LOX inhibition could inhibit matrix stiffness and re-sensitize tumors to therapy (51-54), however LOX inhibition may not be beneficial to treat tumors with existing mature collagen mesh.

Mechanistically, we show that LOX lies upstream of FAK activation; LOX knock-down abolishes basal and TGF-β–induced pFAK (Y397) and collapses focal-adhesion maturation. Activated FAK, in turn, signals through p38 MAPK to inactivate GSK3β at Ser9, allowing β-catenin stabilization and nuclear accumulation. This non-canonical, Wnt-independent β-catenin/YAP signaling regulate CAF differentiation and melanoma progression (37). Our inhibitor data further reveal that Akt provides a permissive background level of GSK3β-Ser9 phosphorylation, but p38—primed by the LOX-FAK axis—is indispensable for full inhibition of GSK3β and robust β-catenin activity.

Clinically, LOX is markedly enriched in the stromal, but not epithelial, compartment of both IDC and DCIS (Oncomine analysis). Functionally, LOX-deficient CAFs lose the capacity to accelerate HCC1806 tumor growth in vivo. Whether this attenuation arises solely from diminished matrix cross-linking or also involves paracrine modulation of immune or cancer cells requires future investigation.

From a therapeutic standpoint, LOX inhibitors (e.g., LXG6403) if exhibit favorable pharmacokinetics could be used for on-target stromal modulation (55). Our work provides a mechanistic rationale for repurposing such agents to disrupt the LOX-FAK-β-catenin axis in desmoplastic breast cancers—potentially in combination with FAK or p38 inhibitors. In summary, we uncover a LOX/FAK/β-catenin cascade that locks mammary fibroblasts into a myCAF phenotype and drives CAF-mediated tumor promotion. Targeting LOX or its downstream effectors could therefore re-program the tumor microenvironment and blunt breast-cancer progression.

### Limitations of the study

The major limitation of this study is that all in vivo experiments relied on xenografts of a single human breast-cancer cell line co-implanted with human CAFs into immunocompromised mice, which lack functional T- and B-cell immunity. Consequently, the role of stromal LOX was assessed only in the context of its biomechanical and paracrine effects on tumor cells; any influence on immune cell recruitment, checkpoint signaling or T cell exclusion—now recognized as critical dimensions of CAF biology—could not be captured. Moreover, pharmacodynamic endpoints such as collagen cross-linking and matrix stiffness may differ in an intact immune microenvironment, limiting the direct translational relevance of our findings. Validation in syngeneic or humanized, immune-competent models will therefore be essential to determine whether targeting the LOX–FAK– β-catenin axis re-programmes CAFs and improves therapeutic response within the full complexity of the tumor microenvironment.

## Funding and acknowledgements

Authors would like to acknowledge Science and Engineering Research Board (EMR/2016/006270 (to PK) and ECR/2015/000280 (to RB)), the Department of Biotechnology (FA/DBTO-18-0005.05), Government of India and the Institute of Eminence grant from the Indian Institute of Science to RB (IE/CARE-19-0319) for financial support. AG acknowledges Council of Scientific and Industrial Research, New Delhi for fellowship (09/079(2778)/2018-EMR-1). PK is a recipient of Fellowship from Indian National Science Academy. RB is a recipient of an intermediate fellowship from the India Alliance DBT Wellcome Trust. Authors are grateful to Sheetal Tushir and Sathisha Kamanna for assistance with online data submission; Quantitative PCR and confocal core facilities at Indian Institute of Science. We acknowledge the use of large language models (e.g., ChatGPT) to assist with grammar refinement for the manuscript text.

## Author contributions

AG and PK conceptualized and designed the study and wrote the paper. AG performed the experiments and analysed the data. SH provided technical help. DP, SM, AR and RB provided human breast cancer tissues for CAFs. DB and UT performed LCI MS/MS analyses. RB, UT, RVK and PK provided the reagents.

## Figure legends

**Figure S1. Comparative secretome analysis of TGF-β over-expressing normal mammary fibroblasts and breast cancer-associated fibroblasts. (A)** Representative image of immunocytochemical analysis for α-SMA and vimentin in CAFs isolated from breast cancer biopsy tissues. α-SMA and vimentin are upregulated in CAFs as compared to HMF3s. Scale bar 10 µm. Magnification 63 ×. **(B)** Semi-quantitative PCR analysis of porcine TGF-β in HMF3s cells. Primers are specific for porcine TGF-β and therefore do not amplify human TGF-β. **(C)** PAI-luciferase reporter assay to detect active TGF-β in the conditioned media of HMF3s with or without TGF-β over-expression. Plot shows RLU (Relative Luciferase Units) which is an indicator of TGF-β induced PAI promoter activity. N = 3, ***p value < 0.001. **(D - G)** Venn diagrams and Reactome pathway analysis (STRING database) of overlaping proteins identified in myCAF-2 and myCAF-3 with respect to HMF3s conditioned media with or without TGF-β over-expression. Twenty proteins in myCAF-2 and Nine proteins in myCAF-3 exclusively overlapping with TGF-β over-expressing HMF3s are depicted in the figures. **(H)** Validation of MS analysis by quantitative PCR in HMF3sTGF-β compared to vector control of genes that were randomly selected from secretome analysis. **(I)** Regulation of genes shown in “H” in HMF3s cells treated with TGF-β. **(J)** Regulation of genes shown in “H” in CAF from five or more different breast cancer patients. Each point depicts fold change of individual CAF wit h respect to HMF3s cells. Line represents median fold change. *P value < 0.05; ** P value < 0.01 ***P value < 0.001

**Figure. S2. TGF-β induced LOX stabilizes β-catenin through GSK3β inactivation during fibroblast activation. (A)** Immunocytochemical analysis of β-catenin and Smad3 in HMF3s cells were treated with TGF-β. TGF-β treatment increased total β-catenin in HMF3s cells. Scale bar 10 µm, magnification 63×. **(B)** Analysis of Wnt2 and Wnt4 transcripts by semi-quantitative PCR in TGF-β treated HMF3s cells at 24 and 48 hours. hfhTERT cDNA was used as positive control. RPL35a is the internal control. **(C)** Expression of ActA2 RNA (α-SMA transcript) in TGF-β treated HMF3s cells upon knockdown of LOX. ActA2 is compromised in LOX knockdown cells.

**Figure S3: LOX sustains the myCAF phenotype and fuels CAF-driven mammary-tumour growth in vivo (A)** Western blot analysis of LOX in CAFs. LOX was knockdown in CAFs using shRNA G4. α-tubulin is the internal control. **(B)** The plot represents HCC1806 tumor volume with respect to time. Tumor volumes are significantly higher for HCC1806 injected with CAF-4a scrambled control which was compromised upon LOX knockdown. **(C)** Representative images of bioluminescence of HCC1806 tumours. The plot represents photon flux of HCC1806 tumours measured weekly normalized to the signal obtained at the first week of injection. HCC1806 showed significantly increased growth when injected with CAF-4a cells which was compromised upon LOX knockdown in CAF-4a. N = 5, *p value < 0.05, ***p value < 0.005.

**Table S1:** List of gene specific primer sequences.

## Bibliography

1. Augsten M. Cancer-associated fibroblasts as another polarized cell type of the tumor microenvironment. Front Oncol. 2014;4:62.

2. Kalluri R. The biology and function of fibroblasts in cancer. Nat Rev Cancer. 2016;16(9):582–98.

3. Wu SZ, Al-Eryani G, Roden DL, Junankar S, Harvey K, Andersson A, et al. A single-cell and spatially resolved atlas of human breast cancers. Nat Genet. 2021;53(9):1334–47.

4. Bartoschek M, Oskolkov N, Bocci M, Lovrot J, Larsson C, Sommarin M, et al. Spatially and functionally distinct subclasses of breast cancer-associated fibroblasts revealed by single cell RNA sequencing. Nat Commun. 2018;9(1):5150.

5. Hinz B, Phan SH, Thannickal VJ, Prunotto M, Desmouliere A, Varga J, et al. Recent developments in myofibroblast biology: paradigms for connective tissue remodeling. Am J Pathol. 2012;180(4):1340–55.

6. Ohlund D, Handly-Santana A, Biffi G, Elyada E, Almeida AS, Ponz-Sarvise M, et al. Distinct populations of inflammatory fibroblasts and myofibroblasts in pancreatic cancer. J Exp Med. 2017;214(3):579–96.

7. Ghazimoradi MH, Babashah S. The transcriptional regulators GATA6 and TET1 regulate the TGF-beta pathway in cancer-associated fibroblasts to promote breast cancer progression. Cell Death Discov. 2025;11(1):164.

8. Liu X, Li J, Yang X, Li X, Kong J, Qi D, et al. Carcinoma-associated fibroblast-derived lysyl oxidase-rich extracellular vesicles mediate collagen crosslinking and promote epithelial-mesenchymal transition via p-FAK/p-paxillin/YAP signaling. Int J Oral Sci. 2023;15(1):32.

9. Takahashi T, Tomonobu N, Kinoshita R, Yamamoto KI, Murata H, Komalasari N, et al. Lysyl oxidase-like 4 promotes the invasiveness of triple-negative breast cancer cells by orchestrating the invasive machinery formed by annexin A2 and S100A11 on the cell surface. Front Oncol. 2024;14:1371342.

10. O’Hare MJ, Bond J, Clarke C, Takeuchi Y, Atherton AJ, Berry C, et al. Conditional immortalization of freshly isolated human mammary fibroblasts and endothelial cells. Proc Natl Acad Sci U S A. 2001;98(2):646–51.

11. Proia DA, Kuperwasser C. Reconstruction of human mammary tissues in a mouse model. Nat Protoc. 2006;1(1):206–14.

12. Sahai E, Astsaturov I, Cukierman E, DeNardo DG, Egeblad M, Evans RM, et al. A framework for advancing our understanding of cancer-associated fibroblasts. Nat Rev Cancer. 2020;20(3):174–86.

13. Samuel SK, Hurta RA, Kondaiah P, Khalil N, Turley EA, Wright JA, et al. Autocrine induction of tumor protease production and invasion by a metallothionein-regulated TGF-beta 1 (Ser223, 225). EMBO J. 1992;11(4):1599–605.

14. Kondaiah P, Van Obberghen-Schilling E, Ludwig RL, Dhar R, Sporn MB, Roberts AB. cDNA cloning of porcine transforming growth factor-beta 1 mRNAs. Evidence for alternate splicing and polyadenylation. J Biol Chem. 1988;263(34):18313–7.

15. Abe M, Harpel JG, Metz CN, Nunes I, Loskutoff DJ, Rifkin DB. An assay for transforming growth factor - beta using cells transfected with a plasminogen activator inhibitor-1 promoter-luciferase construct. Anal Biochem. 1994;216(2):276–84.

16. Arany PR, Nayak RS, Hallikerimath S, Limaye AM, Kale AD, Kondaiah P. Activation of latent TGF-beta1 by low-power laser in vitro correlates with increased TGF-beta1 levels in laser-enhanced oral wound healing. Wound Repair Regen. 2007;15(6):866–74.

17. Ngo P, Ramalingam P, Phillips JA, Furuta GT. Collagen gel contraction assay. Methods Mol Biol. 2006;341:103–9.

18. Singh M, Beri D, Nageshan RK, Chavaan L, Gadara D, Poojary M, et al. A secreted Heat shock protein 90 of Trichomonas vaginalis. PLoS Negl Trop Dis. 2018;12(5):e0006493.

19. Deutsch EW, Bandeira N, Sharma V, Perez-Riverol Y, Carver JJ, Kundu DJ, et al. The ProteomeXchange consortium in 2020: enabling ‘big data’ approaches in proteomics. Nucleic Acids Res. 2020;48(D1):D1145–D52.

20. Perez-Riverol Y, Csordas A, Bai J, Bernal-Llinares M, Hewapathirana S, Kundu DJ, et al. The PRIDE database and related tools and resources in 2019: improving support for quantification data. Nucleic Acids Res. 2019;47(D1):D442–D50.

21. Arina A, Idel C, Hyjek EM, Alegre ML, Wang Y, Bindokas VP, et al. Tumor-associated fibroblasts predominantly come from local and not circulating precursors. Proc Natl Acad Sci U S A. 2016;113(27):7551–6.

22. LeBleu VS, Kalluri R. A peek into cancer-associated fibroblasts: origins, functions and translational impact. Dis Model Mech. 2018;11(4).

23. Kendall RT, Feghali-Bostwick CA. Fibroblasts in fibrosis: novel roles and mediators. Front Pharmacol. 2014;5:123.

24. Kondaiah P, Pant I, Khan I. Molecular pathways regulated by areca nut in the etiopathogenesis of oral submucous fibrosis. Periodontol 2000. 2019;80(1):213–24.

25. Bordignon P, Bottoni G, Xu X, Popescu AS, Truan Z, Guenova E, et al. Dualism of FGF and TGF-beta Signaling in Heterogeneous Cancer-Associated Fibroblast Activation with ETV1 as a Critical Determinant. Cell Rep. 2019;28(9):2358–72 e6.

26. Lohr M, Schmidt C, Ringel J, Kluth M, Muller P, Nizze H, et al. Transforming growth factor-beta1 induces desmoplasia in an experimental model of human pancreatic carcinoma. Cancer Res. 2001;61(2):550–5.

27. Kojima Y, Acar A, Eaton EN, Mellody KT, Scheel C, Ben-Porath I, et al. Autocrine TGF-beta and stromal cell-derived factor-1 (SDF-1) signaling drives the evolution of tumor-promoting mammary stromal myofibroblasts. Proc Natl Acad Sci U S A. 2010;107(46):20009–14.

28. Barker HE, Cox TR, Erler JT. The rationale for targeting the LOX family in cancer. Nat Rev Cancer. 2012;12(8):540–52.

29. Tenti P, Vannucci L. Lysyl oxidases: linking structures and immunity in the tumor microenvironment. Cancer Immunol Immunother. 2020;69(2):223–35.

30. Zeltz C, Pasko E, Cox TR, Navab R, Tsao MS. LOXL1 Is Regulated by Integrin alpha11 and Promotes Non-Small Cell Lung Cancer Tumorigenicity. Cancers (Basel). 2019;11(5).

31. Kirschmann DA, Seftor EA, Fong SF, Nieva DR, Sullivan CM, Edwards EM, et al. A molecular role for lysyl oxidase in breast cancer invasion. Cancer Res. 2002;62(15):4478–83.

32. Chopra S, Kumar N, Rangarajan A, Kondaiah P. Context dependent non canonical WNT signaling mediates activation of fibroblasts by transforming growth factor-beta. Exp Cell Res. 2015;334(2):246–59.

33. Akhmetshina A, Palumbo K, Dees C, Bergmann C, Venalis P, Zerr P, et al. Activation of canonical Wnt signalling is required for TGF-beta-mediated fibrosis. Nat Commun. 2012;3:735.

34. McCubrey JA, Steelman LS, Bertrand FE, Davis NM, Sokolosky M, Abrams SL, et al. GSK-3 as potential target for therapeutic intervention in cancer. Oncotarget. 2014;5(10):2881–911.

35. Ding Q, Xia W, Liu JC, Yang JY, Lee DF, Xia J, et al. Erk associates with and primes GSK-3beta for its inactivation resulting in upregulation of beta-catenin. Mol Cell. 2005;19(2):159–70.

36. Choi CH, Lee BH, Ahn SG, Oh SH. Proteasome inhibition-induced p38 MAPK/ERK signaling regulates autophagy and apoptosis through the dual phosphorylation of glycogen synthase kinase 3beta. Biochem Biophys Res Commun. 2012;418(4):759–64.

37. Liu T, Zhou L, Yang K, Iwasawa K, Kadekaro AL, Takebe T, et al. The beta-catenin/YAP signaling axis is a key regulator of melanoma-associated fibroblasts. Signal Transduct Target Ther. 2019;4:63.

38. Damineni S, Balaji SA, Shettar A, Nayanala S, Kumar N, Kruthika BS, et al. Expression of tripartite motif-containing protein 28 in primary breast carcinoma predicts metastasis and is involved in the stemness, chemoresistance, and tumor growth. Tumour Biol. 2017;39(4):1010428317695919.

39. Hu D, Li Z, Zheng B, Lin X, Pan Y, Gong P, et al. Cancer-associated fibroblasts in breast cancer: Challenges and opportunities. Cancer Commun (Lond). 2022;42(5):401–34.

40. Chitty JL, Yam M, Perryman L, Parker AL, Skhinas JN, Setargew YFI, et al. A first-in-class pan-lysyl oxidase inhibitor impairs stromal remodeling and enhances gemcitabine response and survival in pancreatic cancer. Nat Cancer. 2023;4(9):1326–44.

41. Elyada E, Bolisetty M, Laise P, Flynn WF, Courtois ET, Burkhart RA, et al. Cross-Species Single-Cell Analysis of Pancreatic Ductal Adenocarcinoma Reveals Antigen-Presenting Cancer-Associated Fibroblasts. Cancer Discov. 2019;9(8):1102–23.

42. Zhang F, Ma Y, Li D, Wei J, Chen K, Zhang E, et al. Cancer associated fibroblasts and metabolic reprogramming: unraveling the intricate crosstalk in tumor evolution. J Hematol Oncol. 2024;17(1):80.

43. Cords L, Tietscher S, Anzeneder T, Langwieder C, Rees M, de Souza N, et al. Cancer-associated fibroblast classification in single-cell and spatial proteomics data. Nat Commun. 2023;14(1):4294.

44. Barcellos-Hoff MH, Akhurst RJ. Transforming growth factor-beta in breast cancer: too much, too late. Breast Cancer Res. 2009;11(1):202.

45. Ge J, Jiang H, Chen J, Chen X, Zhang Y, Shi L, et al. TGF-beta signaling orchestrates cancer-associated fibroblasts in the tumor microenvironment of human hepatocellular carcinoma: unveiling insights and clinical significance. BMC Cancer. 2025;25(1):113.

46. Carthy JM. TGFbeta signaling and the control of myofibroblast differentiation: Implications for chronic inflammatory disorders. J Cell Physiol. 2018;233(1):98–106.

47. Cheon DJ, Tong Y, Sim MS, Dering J, Berel D, Cui X, et al. A collagen-remodeling gene signature regulated by TGF-beta signaling is associated with metastasis and poor survival in serous ovarian cancer. Clin Cancer Res. 2014;20(3):711–23.

48. Payne SL, Fogelgren B, Hess AR, Seftor EA, Wiley EL, Fong SF, et al. Lysyl oxidase regulates breast cancer cell migration and adhesion through a hydrogen peroxide-mediated mechanism. Cancer Res. 2005;65(24):11429–36.

49. Taylor MA, Amin JD, Kirschmann DA, Schiemann WP. Lysyl oxidase contributes to mechanotransduction-mediated regulation of transforming growth factor-beta signaling in breast cancer cells. Neoplasia. 2011;13(5):406–18.

50. Li J, Wang X, Liu R, Zi J, Li Y, Li Z, et al. Lysyl Oxidase (LOX) Family Proteins: Key Players in Breast Cancer Occurrence and Progression. J Cancer. 2024;15(16):5230–43.

51. Huang J, Zhang L, Wan D, Zhou L, Zheng S, Lin S, et al. Extracellular matrix and its therapeutic potential for cancer treatment. Signal Transduct Target Ther. 2021;6(1):153.

52. Burchard PR, Ruffolo LI, Ullman NA, Dale BS, Dave YA, Hilty BK, et al. Pan-lysyl oxidase inhibition disrupts fibroinflammatory tumor stroma, rendering cholangiocarcinoma susceptible to chemotherapy. Hepatol Commun. 2024;8(8).

53. Miller BW, Morton JP, Pinese M, Saturno G, Jamieson NB, McGhee E, et al. Targeting the LOX/hypoxia axis reverses many of the features that make pancreatic cancer deadly: inhibition of LOX abrogates metastasis and enhances drug efficacy. EMBO Mol Med. 2015;7(8):1063–76.

54. Kanapathipillai M, Mammoto A, Mammoto T, Kang JH, Jiang E, Ghosh K, et al. Inhibition of mammary tumor growth using lysyl oxidase-targeting nanoparticles to modify extracellular matrix. Nano Lett. 2012;12(6):3213–7.

55. Cetin M, Saatci O, Rezaeian AH, Rao CN, Beneker C, Sreenivas K, et al. A highly potent bi-thiazole inhibitor of LOX rewires collagen architecture and enhances chemoresponse in triple-negative breast cancer. Cell Chem Biol. 2024;31(11):1926–41 e11.

