## Supplementary figures and images for "Stromal LOX–FAK–β-catenin pathway locks mammary fibroblasts into a tumor-promoting myCAF state"

### Supplemental Figure S1

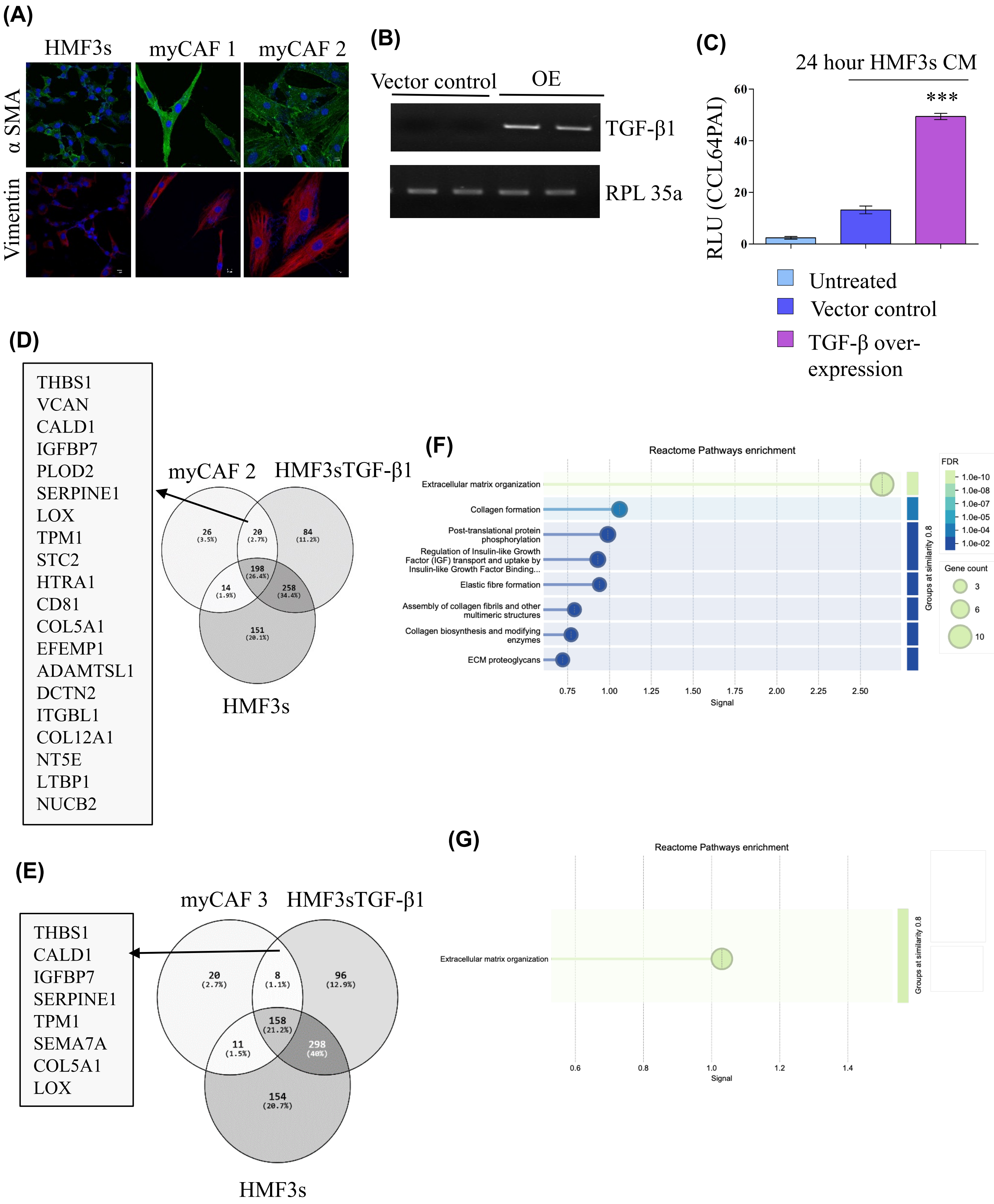

**Figure S1 (Continued)**

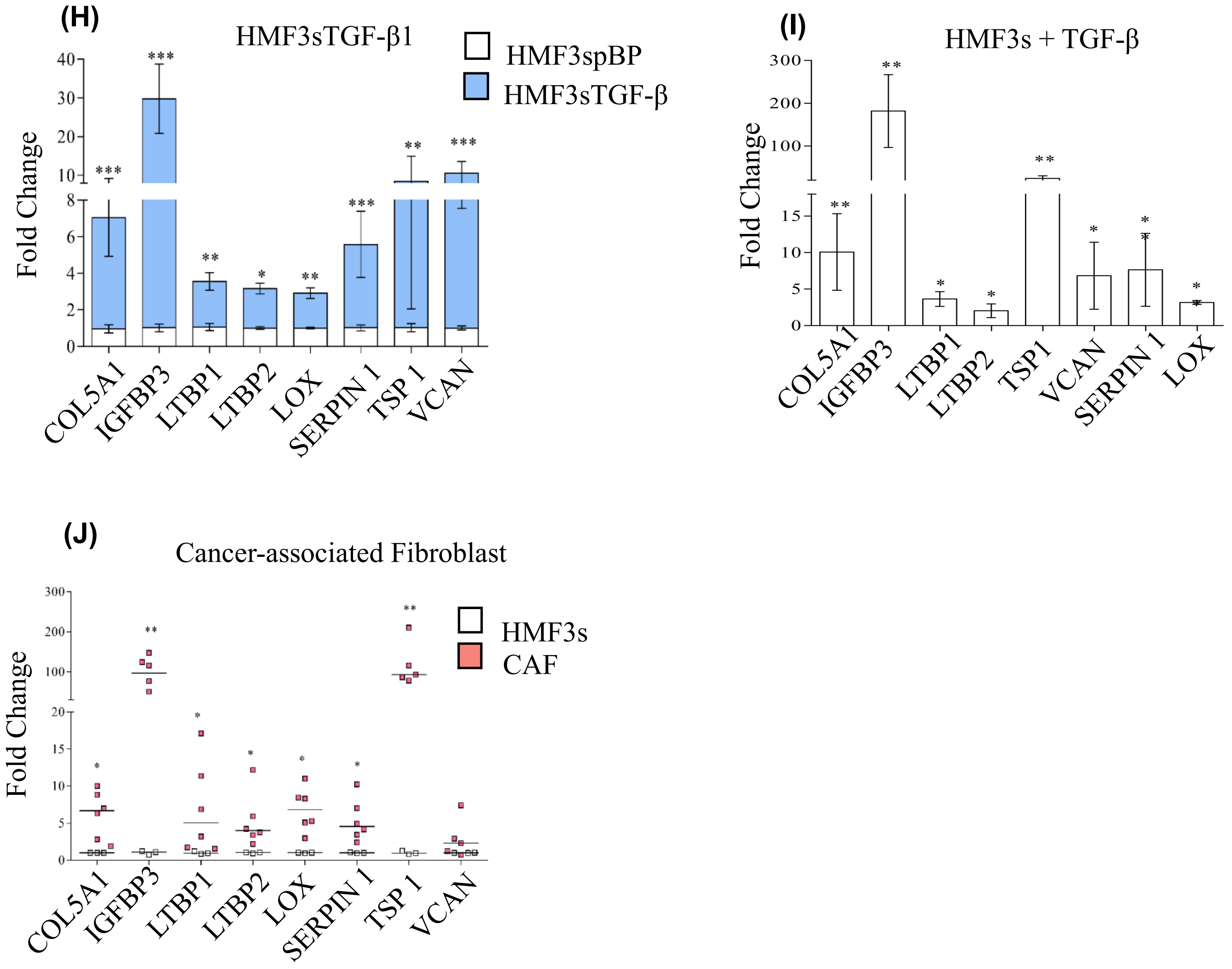

### Supplemental Figure S2

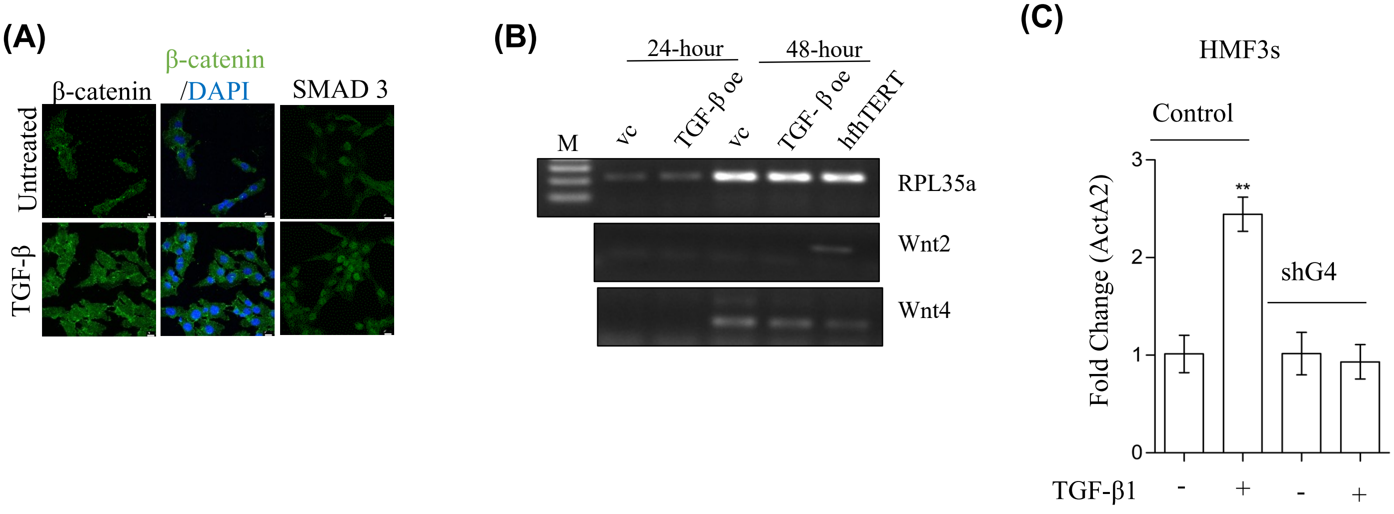

### Supplemental Figure S3

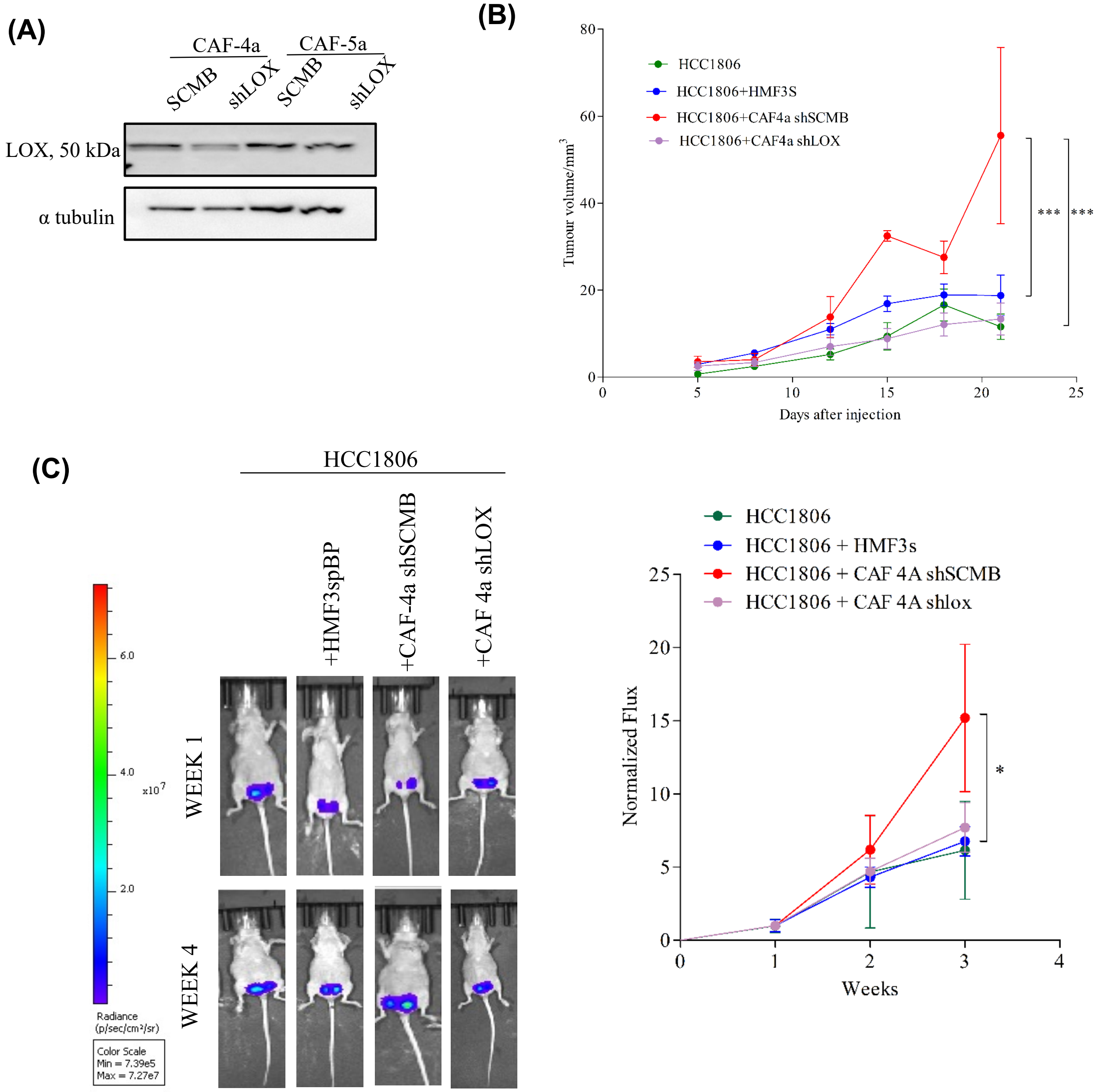
