## Supplemental Table 1 for "Stromal LOX–FAK–β-catenin pathway locks mammary fibroblasts into a tumor-promoting myCAF state"

**Supplementary Table 1: List of gene specific primers used for PCR**

| No. | Gene | Forward primer | Reverse primer |
|---|---|---|---|
| 1 | α-SMA/ActA2 | 5’- CAGCCAAGCACTGTCAGG -3’ | 5’- CAATGGATGGGAAAACAGC -3’ |
| 2 | COL5A1 | 5’- CCATACCCGCTGGAAAGC -3’ | 5’- TCAGGCAAGTTGTGAAAATCT -3’ |
| 3 | IGFBP3 | 5’- AGAGCACAGATACCCAGAACT -3’ | 5’- TGAGGAACTTCAGGTGATTCAGT -3’ |
| 4 | LOX | 5’- GGCTGCTGAAGAAAGCTC -3’ | 5’- CACCACCGGATACTTGGT -3’ |
| 5 | LTBP1 | 5’- AATGATGGAATGCCTACCGGG -3’ | 5’- GTCCTGCTCCACAGATATCA -3’ |
| 6 | LTBP2 | 5’- GGACGCGGAGTGTGTGAATACC -3’ | 5’- GGGTAGCAGAAGCAGCAGCAGTAGG -3’ |
| 7 | Porcine TGF-β1 | 5’- CCGGAACCTGTATTGCTCTC -3’ | 5’- GGCGAAAACCCTCTATAGCC -3’ |
| 8 | RPL 35A | 5’- GAACCAAAGGGAGCACACAG -3’ | 5’- CAATGGCCTTAGCAGGAAGA -3’ |
| 9 | SERPIN 1 | 5’- ATCGAGGTGAACGAGAGTGG -3’ | 5’- AGTGTTCCTGTGGGGTTGTG -3’ |
| 10 | TSP1 | 5’- CCGGCGTGAAGTGTACTAGCTA -3’ | 5’- TGCACTTGGCGTTCTTGTT -3’ |
| 11 | VCAN | 5’- ACAAGCATCCTGTCTCACG -3’ | 5’- TGAAACCATCTTTGCAGTGG -3’ |
